# MvfR shapes *Pseudomonas aeruginosa* Interactions in Polymicrobial Contexts: Implications for Targeted Quorum Sensing Inhibition

**DOI:** 10.1101/2025.05.16.654325

**Authors:** Kelsey M. Wheeler, Myung Whan Oh, Julianna Fusco, Aishlinn Mershon, Erin Kim, Antonia De Oliveira, Laurence G. Rahme

**Author notes:** Correspondence: Laurence G. Rahme; Massachusetts General Hospital 340 Thier Research Building 50 Blossom Street Boston, MA 02114.

## Abstract

Infections often occur in complex niches consisting of multiple bacteria. Despite the in-creasing awareness, there is a fundamental gap in understanding which interactions govern mi-crobial community composition. Pseudomonas aeruginosa is frequently isolated from monomicrobi-al and polymicrobial human infections. This pathogen forms polymicrobial infections with other ESKAPEE pathogens and defies eradication by conventional therapies. By analyzing the competi-tion within cocultures of P. aeruginosa and representative secondary pathogens that commonly co-infect patients, we demonstrate the antagonism of P. aeruginosa against other ESKAPEE pathogens and the contribution of this pathogen’s multiple quorum sensing (QS) systems in these interac-tions. QS is a highly conserved bacterial cell-to-cell communication mechanism that coordinates collective gene expressions at the population level, and it is also involved in P. aeruginosa virulence. Using a collection of P. aeruginosa QS mutants of the three major systems, LasR/LasI, MvfR/PqsABCDE, and RhlR/RhlI and mutants of several QS-regulated functions, we reveal that MvfR and, to a lesser extent, LasR and RhlR control competition between P. aeruginosa and other microbes, possibly through their positive impact on pyoverdine, pyochelin, and phenazine genes. We show that MvfR inhibition alters competitive interspecies interactions and preserves the coexistence of P. aeruginosa with ESKAPEE pathogens tested while disarming the pathogens’ ability to form biofilm and adhere to lung epithelial cells. Our results highlight the role of MvfR inhibition in modulating microbial competitive interactions across multiple species, while simultaneously atten-uating virulence traits. These findings reveal the complexity and importance of QS in interspecies interactions and underscore the impact of the anti-virulence approach in microbial ecology and its importance for treating polymicrobial infections.

## 1. Introduction

Bacteria most often exist within complex microbial communities, where bacteria interact in multiple ways, including the secretion of inhibitory molecules, virulence factor regulation, and biofilm formation [1]. These behaviors facilitate competitive, cooperative, or neutral relationships between bacteria. During infection, these interspecies interactions have the potential to influence pathogen fitness, infection severity, and antibiotic susceptibility.

*Pseudomonas aeruginosa* (PA from here on) is a highly adaptable bacterium that ubiquitously inhabits diverse environments, including soil, marine habitats, plants, and animals [2–5]. PA often establishes polymicrobial communities in human sites, including lung, burn wounds, gut, and environmental reservoirs [1]. This pathogen secretes antimicrobial chemicals and communicates and cooperates with other organisms to develop a multispecies community. For example, PA commonly causes difficult-to-treat polymicrobial infections with other ESKAPEE pathogens (*Enterococcus faecium* (EF from here on), *Staphylococcus aureus* (SA from here on), *Klebsiella pneumoniae* (KP from here on), *Acinetobacter baumannii* (AB from here on), Enterobacter species, and *Escherichia coli* (EC from here on)), which “escape” killing by antibiotics [6–10]. They are particularly concerning since they represent the largest group of nosocomial pathogens. PA is a common dominator in polymicrobial infections due to multiple mechanisms allowing its rapid adaptation to the specific conditions of the host [1].

Quorum sensing (QS) is the central regulatory system that controls key group behaviors such as antibiotic production, biofilm formation, production of virulence factors, expression of bioluminescence, sporulation, and more. Several QS systems in PA have been described to date, and at least three main QS systems are found in PA: the N-acyl-homoserine lactone systems, las (LasR/I) [11] and rhl (RhlR/I) [12, 13], and the MvrR/PqsABCDE system [14–17]. In the host, QS can have a profound impact on the ability of PA to establish infections [18–20]. In particular, the key global virulence transcriptional regulator in PA, MvfR (multiple virulence factor regulator), also referred to as PqsR, controls the synthesis of a range of small molecules, including the 4-hydroxy-2-alkylquinolines (HAQs) [15–17, 21], 2’-aminoacetophenone (2-AA) [22], and 2-n-heptyl-4-hydroxyquinoline N-oxide (HQNO) [23] that promote virulence, competition, and long-term bacterial presence during infection [14, 24–26]. These molecules and methylated alkylquinolones have been reported thus far to be found exclusively in Pseudomonads and Burkholderia species, suggesting they predominantly serve an antagonistic role [27]. By contrast, the LasR and RhlR systems are based on the production of N-acyl homoserine lactone (AHL) signals, a class of structurally conserved molecules that control diverse behaviors, including virulence factor production and biofilm formation in many Gram-negative bacteria [11, 19, 20, 28] and therefore potentially facilitate interspecies cross talk.

The genetic pathways influencing interspecies interactions remain poorly understood. Here, we systematically dissect interactions between PA and other opportunistic pathogens that commonly infect humans, examining the relative importance of exploitative (resource) and interference (antagonistic) competition, two processes thought to play an important role in determining the composition of microbial communities [29]. Moreover, we investigate the impact of an anti-virulence approach by assessing how the presence of an MvfR inhibitor [30] can alter competitive interspecies interactions between pathogens, antibiotic susceptibility for each pathogen, and antagonism of a beneficial microbe. As opposed to antibiotics, anti-virulence drugs are highly selective in their mode of action; they target specific virulence factors in pathogens and do not affect bacterial growth and viability.

Therefore, intuitively, they are expected to have a much lower impact on the gut microbiome than antibiotics that kill the bacteria; a promising concept in the context of many human pathologies in which patients already have disturbed microbiota.

Altogether, these findings provide critical insights into how clinically relevant pairs of microbes may interact and how these interactions may be influenced by therapeutic approaches, with clear implications for treating polymicrobial infections.

## 2. Methods

### 2.1. Strains and growth conditions

Experiments with *Pseudomonas aeruginosa* were conducted using the PA14 (Rahme lab- burn isolate) and isogenic single, double, and triple deletion mutants of the quorum-sensing regulators (lasR, rhlR and mvfR) [31, 32] were provided by Dr. Deborah Hogan. The lasR/rhlR and lasR/rhlR/mvfR, and pqsA deletion strains with MvfR complementation were obtained by electroporation as described previously [33] with the pDN18-MvfR plasmid [16]. Competing secondary pathogen strains included Escherichia coli LF82 (provided by Dr. Wendy Garrett), a clinical isolate of *Klebsiella pneumoniae* (Rahme Lab-burn isolate), *Klebsiella pneumoniae* (ATTC 43816), *Acinetobacter baumannii* 17978 (Dr. Alan Hauser), *Staphylococcus aureus* USA300 (provided by Dr. Gerald Pier), *Staphylococcus epidermidis* 1457 (provided by Dr. Gerald Pier), and a clinical isolate of *Enterococcus faecalis* (Rahme Lab- burn isolate). The mvfR/pDN18pqsABCDE, a strain lacking mvfR with constitutive expression of pqs operon, was constructed by cloning into pDN18 plasmid for electroporation into mvfR mutant strain [34]. The pqsA [15, 17, 35], pqsBC[25, 36], pqsE[17], pqsL[37], and pqsH[15] mutants were constructed as previously reported.

All strains were streaked out from frozen 20% glycerol stocks on Lysogeny Broth (LB) agar plates, except for EF, which was cultured on De Man, Rogosa and Sharpe (MRS) agar plates. All liquid overnight cultures were inoculated from single colonies and grown at 37 °C for monoclonal expansion. PA, EC, *Staphylococcus epidermidis*, SA, KP, and AB were cultured shaking at 180 rpm in 5 mL LB. EF was cultured shaking at 180 rpm in 5 mL MRS broth.

### 2.2. Monoculture and co-culture assays

For all competition experiments, overnight cultures were inoculated 1:1 into fresh Eagle’s Minimum Essential Medium (EMEM, Lonza) in 96-well plates to a final volume of 200 μL and a starting cell density of 0.5–1L×L10L CFU/mL for each strain. Cultures were incubated at 37 °C for 24 h. Following incubation, bacterial densities were measured by selective or differential plating. Cultures were serially (10-fold) diluted in 100 µL PBS, and each dilution was spotted onto selective agar plates to enumerate colony-forming units (CFUs) of each species in co-culture.

PA CFUs were enumerated by plating on Pseudomonas Isolation Agar (PIA), which selectively supports PA growth. EC and KP CFUs were enumerated by plating on LB agar supplemented with 80 μg/mL ampicillin, as the isolates tested are intrinsically resistant to ampicillin at this concentration, while PA growth is inhibited, allowing for selective enumeration. EF CFUs were enumerated by plating on MRS agar. SE or SA CFUs were enumerated by plating on high salt TSA (tryptic soy agar + 7% NaCl). AB was plated on LB agar, and colonies were distinguished from PA based on their distinct morphology (Fig. S1). In all cases, PA CFUs were enumerated directly from PIA, not inferred by subtraction from total counts on LB agar.

If no colonies were observed at the lowest dilution, the CFU/mL was recorded as the detection limit, 1–2L×L10³ CFU/mL.

### 2.3. Crystal violet staining for biofilm quantification

Biofilm formation was quantified using a crystal violet staining assay with modifications [38–40]. The bacterial cultures were streaked on Luria broth (LB) agar for overnight growth at 37°C. The next morning, a single colony was inoculated into LB broth for monoclonal expansion at 37oC with shaking at 200 rpm. Each strain was cultured until OD600= 0.1 in LB broth, followed by 1:1000 dilution after washing three times with PBS. Specifically, then, 200 μL of the diluted culture was added to each well of a 96-well polystyrene microtiter plate (Corning, USA) and incubated statically at 37°C for 16 hours with or without dimethyl sulfoxide (DMSO) vehicle, or D88. After incubation, unbound, planktonic bacteria were gently removed by inverting and tapping, followed by washing with sterile PBS. After the washing steps, the plates were dried for 30 minutes at room temperature. The biofilm biomasses were stained with 0.1% (w/v) crystal violet solution for 30 minutes at room temperature. After staining, excess stain was removed by washing with PBS, and plates were dried for 30 minutes at room temperature. The crystal violet stains in the biofilms were allowed to solubilize with Acetone:Ethanol (2:8) solution for 30 minutes at room temperature. The biofilm production was quantified by measuring the absorbance at 570 nm by the microplate reader (Infinite M Plex, Tecan, Männedorf, Switzerland). Sterile LB was used as a negative control. All biofilm assays were done in triplicate and repeated at least three times independently.

### 2.4. Cell adherence assay

A549 human alveolar epithelial cells (ATCC CCL-185) were cultured in Minimum Essential Medium (MEM) supplemented with 10% fetal bovine serum (FBS) and 1% penicillin-streptomycin at 37°C in 5% CO2. Cells were seeded in 12-well plates at a density of 5 × 105 cells/well and grown to 90% confluency level. Prior to infection, A549 cells were washed thoroughly with PBS and incubated in a serum-free, antibiotics-free MEM for 2 hours. The cells were infected with desired bacteria at a multiplicity of infection (MOI) of 25:1 with or without treatments. After inoculation, 12- well plates were centrifuged at 130 xg for 5 minutes to bring cells in contact with bacteria. Cells were incubated for 1.5 h at 37oC in 5% CO2 for adherence. After incubation, the wells were washed gently three times with PBS to remove non-adherent bacteria. Cells were treated with 0.1% Triton X-100 in PBS for 30 minutes at room temperature, and the lysates were serially diluted for CFU enumeration. All cell adherence assays were conducted in triplicate and repeated at least three times independently.

### 2.5. Data analysis and statistics

We performed a secondary analysis of patient clinical data from the Glue Grant [41]. Permission to use de-identified data was obtained from the Massachusetts General Hospital Institutional Review Board (MGH IRB protocol 2008-P000629/1). Among the 2,002 patients in the dataset, our inclusion criteria identified 166 adult patients (age ≥ 18 years) who suffered burn wounds with infection. This Glue Grant dataset was then sorted in R 3.6.1 using the dyplr package into four categories: 1) patients who are only ever infected by a single microbe type across time (monomicrobial infection), 2) patients who develop multiple infections over time with different microbes (multiple types of monomicrobial infections across time), 3) patients who develop infections with distinct infecting microbes at different body sites on a single day (multiple types of monomicrobial infection across body sites), and 4) patients who develop at least one infection with multiple infecting microbes at a single body site on a single day (polymicrobial infection). A decision tree is presented in Fig. S2A. For each polymicrobial infection event (i.e., body site infected with multiple organisms on a single day), the number of infecting organisms, the type of organisms, and the co-occurring pairs were counted. The infecting organism types were tabulated for each monomicrobial infection event (i.e., body site infected with a single microorganism on a single day). Data are presented in Table S1.

For bacterial competition assays, median log-transformed CFU data are plotted (horizontal line) with each replicate overlayed. Significant differences in CFU data were assessed by two-way ANOVA, with either Šídák’s or Dunnett’s post-test to adjust for multiple comparisons as specified in the supplementary Tables S2-8, which contain the mean log-transformed fold changes in CFU, 95% confidence interval of fold change, and adjusted P-values for each plot. Šídák’s post-test was applied for multiple pairwise comparisons, while Dunnett’s test was used when comparing multiple treatments to a single control. Data were plotted and significance assessed in Prism 10.

## 3. Results

### 3.1 *P. aeruginosa* is a frequent common denominator in human polymicrobial infections

The presence of multiple infection-causing bacteria can complicate treatment strategies due to intrinsic differences in antibiotic susceptibility and polymicrobial interactions that enhance phenotypic and genotypic resistance through QS-dependent interactions such as public goods production (e.g., beta-lactamase or biofilm) or 2-AA triggered persistence. Thus, determining the incidence of polymicrobial infections and identifying common interacting pairs of pathogens is a critical first step in designing effective antimicrobial treatment plans. The relevance of polymicrobial infections has gained attention, particularly in cystic fibrosis and periodontal disease. Here, we use burn wounds as a clinically-relevant niche to investigate the incidence and etiology of polymicrobial infections, as these injuries are prone to infection due to a breach in the protective barriers.

We performed a secondary analysis of clinical data collected from individuals enrolled in the multi-center Inflammation and Host Response to Injury (“Glue Grant”) cohort [41], a longitudinal study to study host changes over time in response to trauma and burn. Of 573 patients who suffered burn injuries, we identified 166 adults (age ≥ 18) who developed at least one burn wound infection. In total, these patients experienced 545 infection events (i.e., presence of ≥ 1 pathogen/burn location/day), with a median [IQR] of 2 [1–4] infection events per patient.

Among the 166 burn patients who developed an infection at the site of injury, 54 patients were only ever infected by a single microbe type across time (Category 1. Monomicrobial infection), 15 patients developed multiple infections over time involving different types of microbes (Category 2. Multiple types of monomicrobial infections across time), 7 patients developed infections with distinct infecting microbes at different body sites on a single day (Category 3. Multiple types of monomicrobial infection across body sites), and 90 patients developed at least one infection with multiple infecting microbes at a single body site on a single day (Category 4. Polymicrobial infection) (Fig. S2B). In total, monomicrobial infection events accounted for 61.1% (333/545) of infection events while polymicrobial infections accounted for 38.9% (212/545) (Fig. S2C). Among the 212 polymicrobial infection events, 68.4% consisted of 2 different microbes, 22.6% consisted of 3 different microbes, 8.0% consisted of 4 microbes, and 0.9% consisted of 5 microbes.

Across the 212 polymicrobial infection events, PA had the highest incidence (occurring in 40.6% of events) (Fig. 1A). During PA polymicrobial infections, the most commonly co-occurring bacterial genus was Staphylococcus, with 14.2% of polymicrobial infections consisting of PA and coagulase-negative Staphylococcus (10.8%) or SA (3.3%). In total, 19.3% of polymicrobial infections involved PA and a Gram-positive, 17.0% involved PA and a Gram-negative, such as KP, Acinetobacter, and E.coli, and 10.4% involved PA and a fungal pathogen (Fig. 1B). These findings highlight PA as a common denominator in polymicrobial burn wound infections.

**Figure 1.**
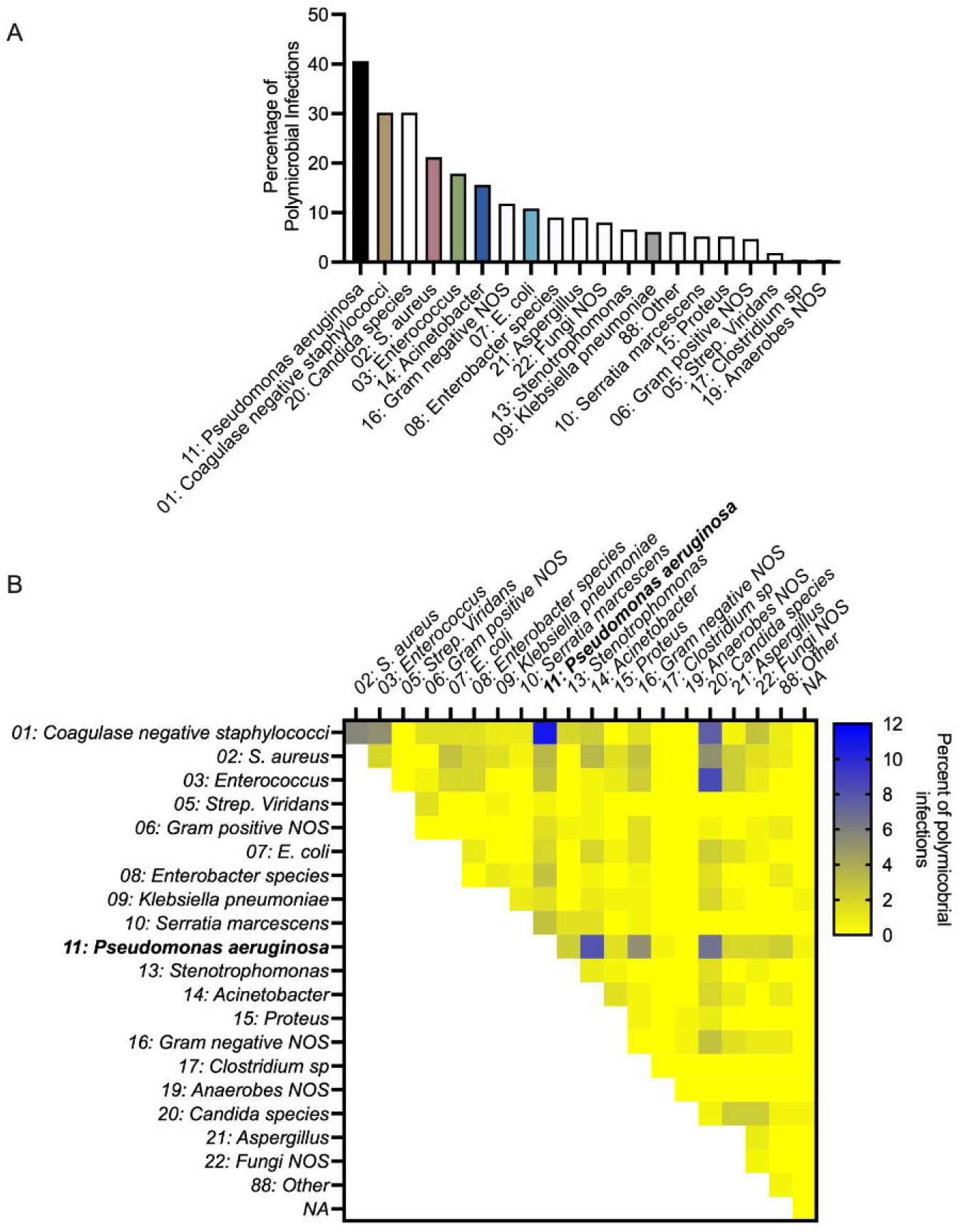
*P. aeruginosa* is common in polymicrobial infections of human burn wounds. A) Percentage of polymicrobial infections consisting of a particular microbe, n = 212 polymicrobial infection events. The quantification of microbes present in monomicrobial or polymicrobial infection events is presented in Table S1. B) Percentage of polymicrobial infections consisting of a pair of microbes, n = 212 polymicrobial infection events.

### 3.2. PA inhibits the growth of Gram-negative ESKAPEE pathogens

Characterizing the interactions within pathogenic polymicrobial communities is important for understanding which pathogens likely suppress the survival of other potential pathogens and which pairs carve out distinct niches to infect the host simultaneously. To this end, we applied a dual-species in vitro competition assay to test whether any growth-inhibiting or growth-promoting interactions can be observed between the bacterial strains common in polymicrobial infections within the Glue Grant dataset. In particular, we focused on interactions between PA strain PA14 and representative bacterial pathogen isolates (EC LF82, Staphylococcus epidermidis 1457 (SE from here on), SA USA300, KP, AB 17897, and EF).

All strains reached similar densities when grown as monocultures in EMEM, and none of the competing strains negatively impacted PA growth (Fig. 2A-F, Table S2). By contrast, KP (Fig. 2A), AB (Fig. 2B), and EC (Fig. 2C) each had several log fewer viable cells when cultured for 24 h in the presence of PA (Table S2). SE had a small < 1-log reduction (Fig. 2E), while SA and EF each had a small < 1-log increase (Fig. 2D, 2F) in biomass in the presence of PA after 24 h (Table S2). EF and PA appeared to engage in a cooperative interaction, with each species reaching a higher density in coculture than in monoculture (Fig. 2F, Table S2).

**Figure 2.**
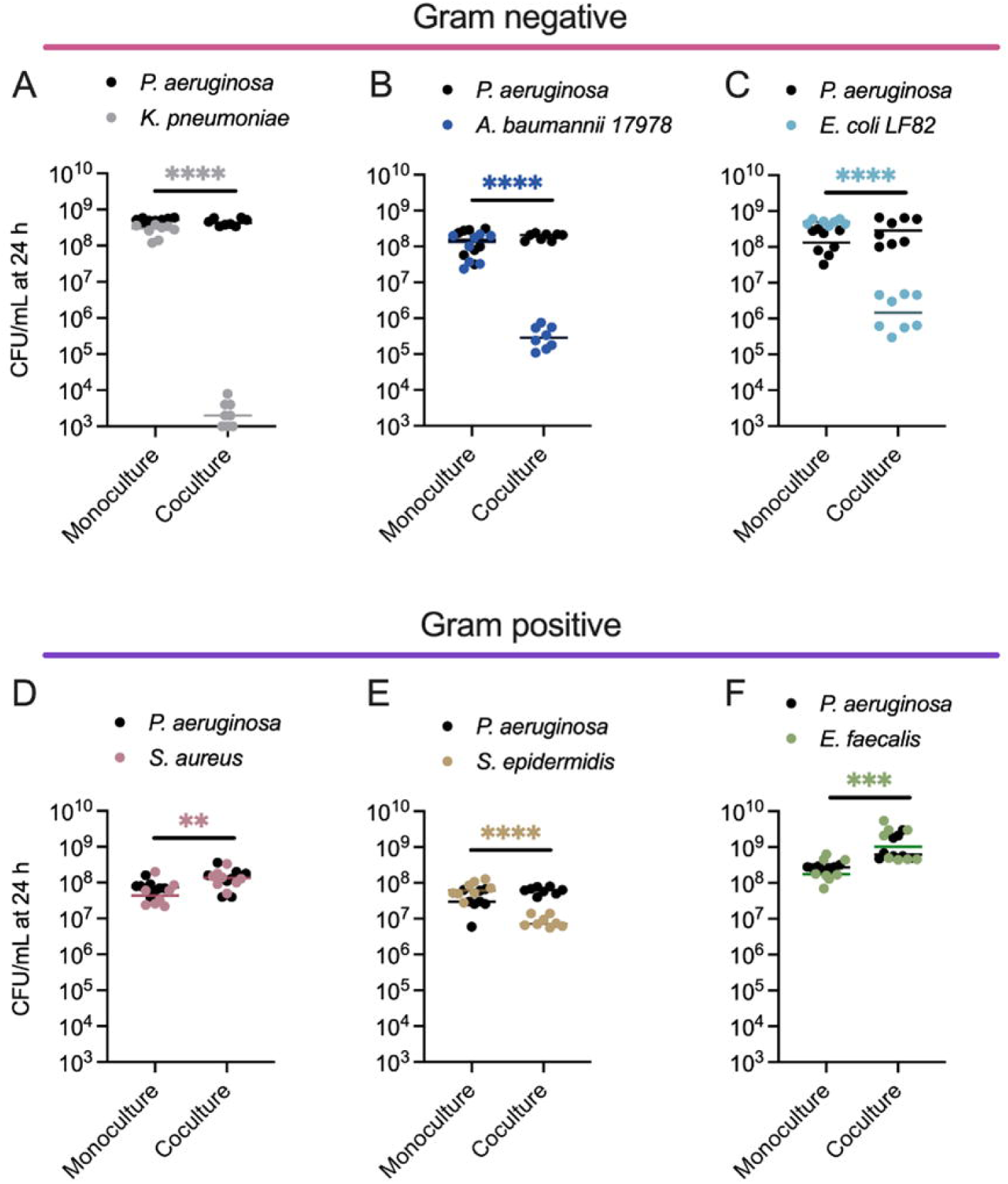
Dual species competitions between P. aeruginosa and other pathogens that co-infect human tissues reveal that PA inhibits the growth of Gram-negative ESKAPEE pathogens. Enumeration of PA and A) KP, B) AB, C) EC, D) SA, E) SE, or F) EF colony forming units (CFU) following 24 h growth in monoculture or co-culture. PA CFU for each replicate is indicated with black circles, and the competing bacterium CFU for each replicate is indicated with different colored circles, n = 8 biological replicates. The middle line indicates the median log-transformed CFU/mL per bacterium in monoculture or co-culture. Significant differences between the mono- vs. co-cultures were assessed by two-way ANOVA on the log10-transformed CFU/mL data, followed by Šídák’s multiple comparisons test. ** P< 0.01, *** P<0.001, **** P<0.0001. Complete statistical analysis is presented in Table S2.

### 3.3. Anti-virulence therapy targeting the quorum sensing regulator, MvfR, restricts competitive interactions between PA and Gram-negative ESKAPEE pathogens

Targeting bacterial virulence with QS inhibitors is an attractive therapeutic strategy for combating infection, as it disarms the pathogen without affecting bacterial growth or viability. The competitive targeting of QS regulators by these compounds against the native signal molecules inhibits the activation of QS signaling, preventing the expressions of downstream genes encoding for virulence factors and pathogenesis [30, 42, 43]. Thus, these compounds are not expected to impose a strong selective pressure on pathogenic bacteria and, in turn, are less likely to promote the evolution of resistant strains. One potent anti-virulence agent we developed recently, a novel N-Aryl Malonamide (D88), is highly soluble, bears no potential substituents incompatible with in vivo use, and directly targets MvfR [30]. While this compound is highly efficacious in the monomicrobial PA infection of burn wounds in mice [30], it is unknown how this QS inhibitor might alter the composition of a polymicrobial community.

To begin addressing this gap, we measured in vitro composition of a two-member community consisting of PA after 24 h in the presence of increasing concentrations of D88, the MvfR-inhibitor. This revealed that disrupting MvfR activity with micromolar concentrations of the D88, but not its vehicle (DMSO), significantly alters the structure of the two-member community (Fig. 3A-C, Table S3), without directly inhibiting the growth of any of these strains even at a concentration exceeding those used in the competition experiments (Fig. S3, S4). The competition between KP, AB, or EC against PA ΔmvfR led to a significant rescue of survival in the secondary pathogens. The large shifts in the distribution of the two pathogens confirmed that the antagonism of PA towards these Gram-negative pathogens involves MvfR, and suggests that MvfR-regulated genes confer a conserved role in bacterial competition with diverse Gram-negative bacteria.

**Figure 3.**
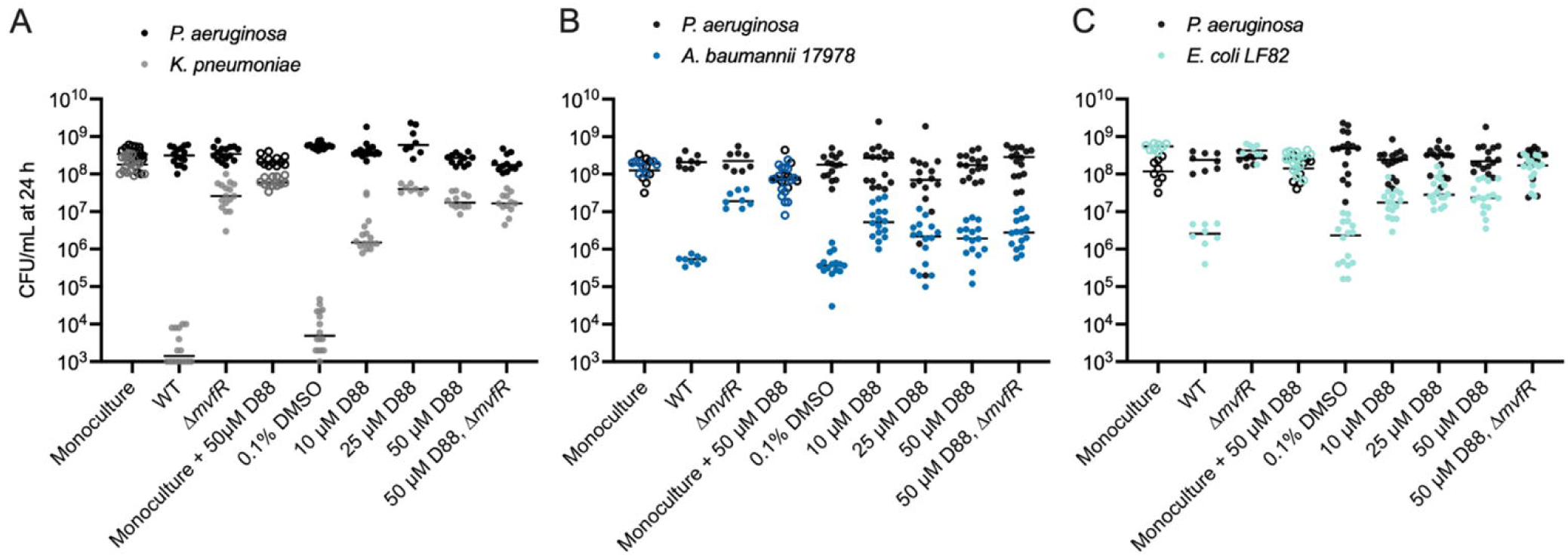
Inhibiting the quorum sensing regulator MvfR alters competitive interactions between PA and Gram-negative ESKAPEE pathogens. A) Enumeration of KP CFU (grey) and PA CFU (black) following 24 h monoculture growth with 50 µM D88 or co-culture with PA WT and increasing concentrations of the MvfR inhibitor D88 or the vehicle alone (DMSO, 0 µM D88). CFUs for monoculture experiments are indicated with open circles, while CFUs for co-culture experiments are indicated with closed circles. n = 8-16 independent replicates. The middle line indicates the median log-transformed CFU/mL of KP or PA in monoculture or co-culture. PA CFU was not strongly altered (< 1-log) by co-culture with KP. B) Enumeration of PA and AB CFU following 24 h mono- and co-culture with or without increasing concentrations of the MvfR inhibitor D88 or the vehicle alone (DMSO). PA WT inhibits AB proliferation compared to monoculture. Deletion of mvfR in PA restores AB viability in co-culture. Low D88 concentrations partially restore the viability of AB in co-culture. PA CFU for each replicate is indicated with black circles, and AB CFU for each replicate is indicated with dark blue circles. CFUs for monoculture experiments are indicated with open circles, while CFUs for co-culture experiments are indicated with closed circles. n = 8-16 independent replicates. C) Enumeration of PA and EC CFU following 24 h mono- and co-culture with or without increasing concentrations of the MvfR inhibitor D88 or the vehicle alone (DMSO). PA WT inhibits EC proliferation compared to monoculture. Deletion of mvfR in PA restores EC viability in co-culture. D88 partially restores EC viability in co-culture. PA CFU for each replicate is indicated with black circles, and EC CFU for each replicate is indicated with cyan circles. CFUs for monoculture experiments are indicated with open circles, while CFUs for co-culture experiments are indicated with closed circles. n = 8-16 independent replicates. For each panel, the middle line indicates the median log-transformed CFU/mL per bacterium in monoculture or co-culture. Significant differences between the mono- vs. co-cultures were assessed by two-way ANOVA on the log10-transformed CFU/mL data, followed by Dunnett’s multiple comparisons test. Statistical analysis is presented in Table S3.

Intriguingly, we find that disrupting MvfR activity with D88 partially rescues the survival of AB in coculture with PA at 10 and 25 µM concentrations without impacting PA abundance (Fig. 3B, Table S3). However, at the concentration of 50 µM D88, AB survival in co-culture is not significantly higher than in the medium alone (Fig. 3B, Table S3). Since AB growth is not inhibited by 50 µM D88 in monoculture (Fig. 3B, Table S3), we hypothesized that a high concentration of D88 may sensitize AB to killing by PA in an MvfR-independent manner. Consistent with this hypothesis, while deletion of mvfR restores AB survival without D88, it does not impact AB survival in co-culture with 50 µM D88 (Fig. 3B, Table S3).

In the case of EC-PA co-cultures, disrupting MvfR activity with micromolar concentrations of the MvfR-inhibitor D88 partially recovered EC survival without impacting PA growth (Fig. 3C, Table S3). D88 did not affect EC viability in monoculture. The deletion of mvfR gene from PA further recovered EC survival in co-cultures with 50 µM D88 (Fig. 3C, Table S3). Taken together, these results highlight the role of MvfR inhibition in tuning microbial competitive interactions in multiple species and, moreover, hint at the possible existence of homologous targets in AB that sensitize it to MvfR-independent antagonism.

### 3.4. MvfR controls interference competition between PA and KP in co-culture

To elucidate the kinetics of PA competition with co-infecting Gram-negative pathogens, we focused on PA-KP dual cultures, which exhibited the most robust competition (i.e., largest change in CFU) among the tested pairs (Fig. 2), providing a sufficient dynamic range to assess changes in competition. A time course of PA and KP growth revealed that both strains exhibit similar growth dynamics in monoculture (Fig. 4A). When cultured together, both strains initially grow to similar densities, reaching a density of approximately 108 cells/mL after 4 h (Fig. 4B). Beyond that point, however, the number of viable KP cells begins to drop significantly by about 5- log over 48 h, while PA density remains relatively constant (Fig. 4B). The greatest variability in CFU counts occurred at 16 h, coinciding with the steepest decline in KP viability. This pattern of rapid exclusion of KP from the culture is most consistent with a mechanism of interference competition, reviewed in [29].

**Figure 4.**
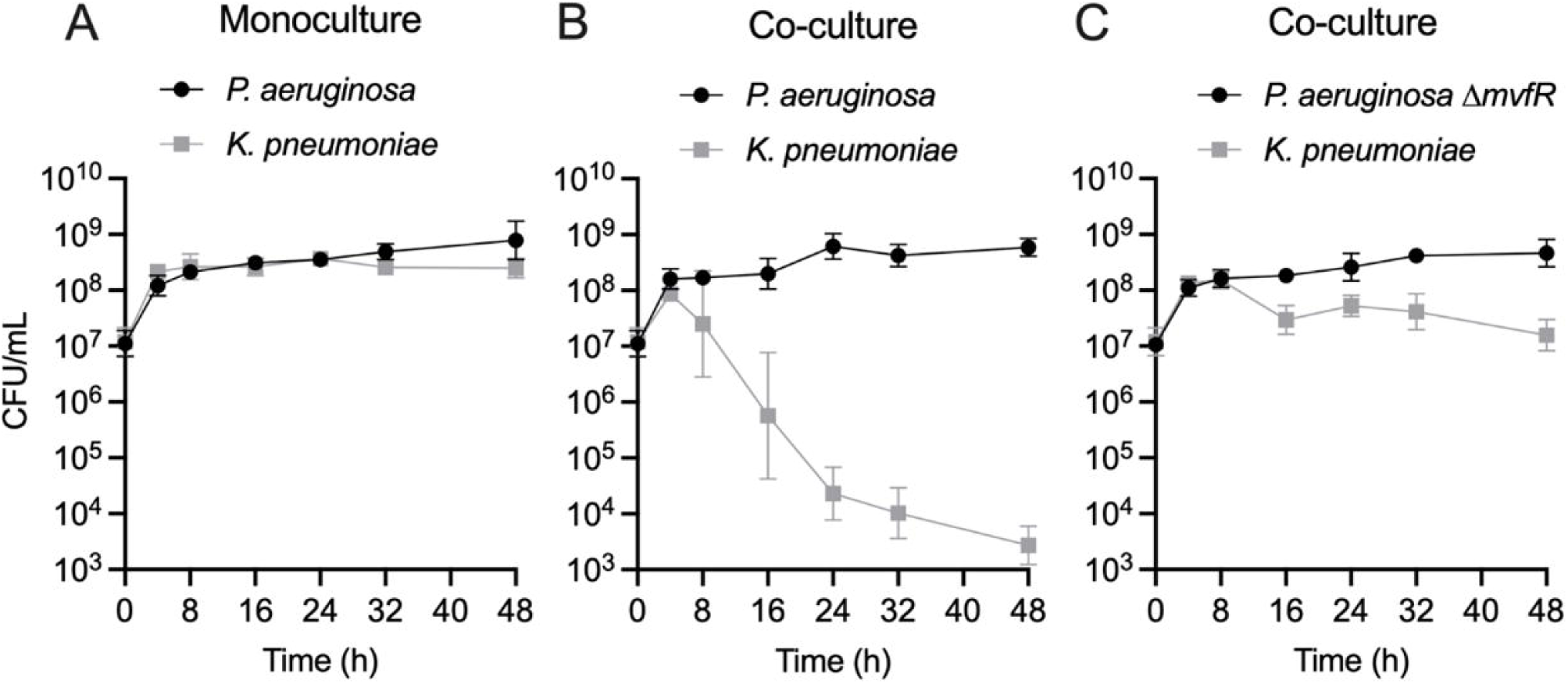
The quorum sensing regulator MvfR is essential for direct antagonism of KP by PA in co-culture. A) Enumeration of PA (Black circle) and KP (Grey square) CFU/mL over 48 h monoculture growth. Data point indicates the mean log-transformed CFU/mL, error bars indicate the 95% confidence interval. n = 8 biological replicates per time point. B) Enumeration of PA (Black circle) and KP (Grey square) CFU/mL over 48 h co-culture growth. Data point indicates the mean log-transformed CFU/mL, error bars indicate the 95% confidence interval. n = 8 biological replicates per time point. C) Enumeration of PA ΔmvfR (Black circle) and KP (Grey square) CFU/mL over 48 h co-culture growth. Data point indicates the mean log-transformed CFU/mL, error bars indicate the 95% confidence interval. n = 8 biological replicates per time point. Inoculating CFU at 0 h was 0.5-1 × 107 CFU/mL.

We further tested whether another clinical isolate of PA obtained from a polymicrobial burn wound infection (PA416A) engages in similar interactions with KP as the PA14 strain (Fig. S5A). Indeed, as with the PA14 strain, we find that the PA416A strain competitively inhibits the growth of KP. Likewise, we find a different competing strain of KP (ATCC 43816) was inhibited by PA14 by 2-log (Fig. S5B).

Importantly, the growth kinetics of KP with PA ΔmvfR suggests that MvfR is essential for the competition observed in EMEM, showing only a small reduction of KP biomass by ∼1-log between 16-48 h (Fig. 4C). This growth reduction/inhibition may arise from the competition for scarce resources as the community depletes nutrients from the medium. However, the MvfR-dependent competition between PA and KP is also evident in a nutrient-rich medium (LB) (Fig. S6), suggesting that these two species may engage in antagonistic interactions in a variety of environments.

### 3.5. The quorum sensing regulator MvfR is a crucial driver of competition between PA and KP

The quorum sensing (QS) system is a density-dependent signaling mechanism that controls numerous behaviors associated with microbial antagonism, including the production of proteases, iron scavenging molecules, and small antimicrobial molecules [19, 44] (Fig. 5A). The three major QS systems in PA, las (LasR/I), rhl (RhlI/R), and mvfR (MvfR/PqsABCDE)) systems, are regulated through an interconnected network of growth phase-dependent interactions whereby LasR, directly and indirectly, activates MvfR and RhlR [45–47], MvfR exerts direct control of the RhlR and LasR systems [44], and RhlR can inhibit MvfR [48] (Fig. 5A). To investigate the contribution of each of these regulators in interspecies competition in vitro, we tested whether PA QS mutant strains (ΔlasR, ΔrhlR, ΔmvfR, ΔlasRΔrhlR, ΔlasRΔmvfR, ΔrhlRΔmvfR, ΔlasRΔrhlRΔmvfR) have a diminished ability to compete with KP. Indeed, extensive gene deletion analyses revealed that these major PA QS systems differentially impact the composition of a two-member community consisting of PA and KP (Fig. 5B, Table S4). Specifically, the deletion of lasR or rhlR increases KP survival by approximately 2-log in co-culture (Fig. 5B). Importantly, however, the deletion of mvfR demonstrated the most prominent effect on KP survival, restoring survival by approximately 4-log without affecting PA growth relative to the wild-type (WT) strain (Fig. 5B, Table S4). Likewise, the QS double and triple mutants (ΔlasRΔmvfR, ΔrhlRΔmvfR, ΔlasRΔrhlR, ΔlasRΔrhlRΔmvfR) all rescued KP survival by approximately 4-log. To investigate whether mvfR expression is sufficient to drive competition between PA and KP independent of the other QS systems, we complemented the ΔlasRΔrhlR and the ΔlasRΔrhlRΔmvfR mutants with a plasmid that constitutively expresses MvfR (pDN18-mvfR) to bypass potential regulatory effects by the Las/Rhl systems and ensure consistent expression. This approach allows us to decouple the impact of LasR and RhlR from MvfR. Complementation of the ΔlasRΔrhlR and the ΔlasRΔrhlRΔmvfR mutant strains restored the inhibitory action of PA against KP by 2-log, or approximately to the level of killing of single lasR or rhlR deletion (Fig. 5B), further highlighting the importance of MvfR in interspecies competition.

**Figure 5.**
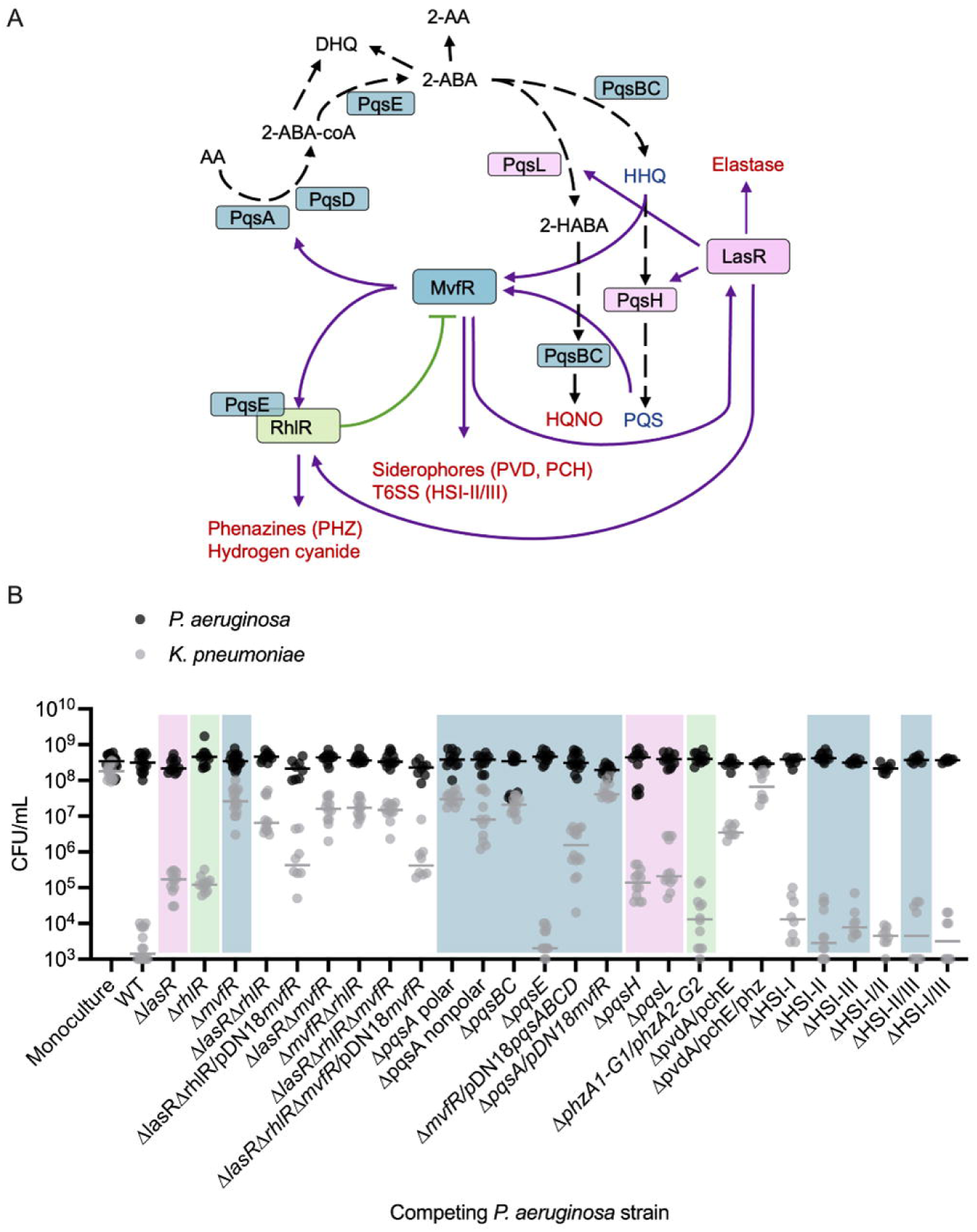
The major quorum sensing systems in PA each contribute to inhibitory interactions with KP in coculture. A) Current view of the QS regulatory network in PA and a subset of its virulence-regulated functions (red text). MvfR, in the presence of PQS or HHQ, binds and activates the transcription of genes, including lasR and rhlR, and the pqsABCD operon and pqsE, whose encoded proteins catalyze the biosynthesis of compounds, including PQS, HHQ, HQNO, and 2-AA. Purple line indicates activation, and green line indicates inhibition, black dashed line indicates biosynthetic process. B) Enumeration of KP CFU (grey) and PA CFU (black) following 24 h mono- or co-culture with different strains of PA. PA WT inhibits KP proliferation in EMEM (n = 16 independent replicates), compared to monoculture (n = 16 independent replicates). Deletion of each of QS regulators (LasR, RhlR, or MvfR) in PA restores KP survival in co-culture, with deletion of mvfR (n = 16 independent replicates) having the strongest effect. Double and triple mutations in any combination of the QS regulators (n = 12 independent replicates) similarly restores KP survival. MvfR complementation (n = 8 independent replicates) of the triple QS mutant and the double LasR/RhlR mutant restores PA antagonism to the level of lasR and rhlR single mutants (n = 12 independent replicates). PA CFU were not significantly altered relative to monoculture in any competition. Colored shaded boxes indicate the primary QS regulator associated with the gene that was mutated: pink for LasR-regulated genes, green for RhlR-regulated genes, and light blue for MvfR-regulated genes. The middle line indicates the median log-transformed CFU/mL of KP or PA in monoculture or co-culture. Significant differences between the mono- vs. co-cultures were assessed by two-way ANOVA on the log10- transformed CFU/mL data, followed by Dunnett’s multiple comparisons test. Complete statistical analysis is presented in Table S4. Note: CFU measurements from Fig. 2A for the monoculture, WT, and ΔmvfR columns are included for comparison as they were completed as a part of the same set of experiments.

We observed the largest variability in CFU counts in conditions where KP survival was intermediate, meaning KP was neither fully eliminated nor completely sustained in the competition. This variability likely reflects dynamic fluctuations in competition dynamics, with factors such as quorum sensing and growth phase influencing the outcome of these interactions.

Through the transcriptional activation of the pqsABCDE operon, MvfR drives the production of its autoinducers, HHQ and PQS, as well as dozens of other small molecules, including the antagonistic factor HQNO [24, 49]. To investigate the involvement of MvfR-regulated functions in interspecies competition, we examined whether deletion mutants lacking key enzymes in the PQS and HQNO biosynthesis pathways (pqsA, pqsBC, pqsH, pqsL) inhibit KP growth. The pqsABCDE operon, under MvfR control, is responsible for HHQ biosynthesis (Fig. 5A), while pqsH and pqsL, which are activated by LasR, control PQS and HQNO synthesis, respectively (Fig. 4A). Co-culturing KP with PA ΔpqsA polar, ΔpqsA nonpolar, or ΔpqsBC nonpolar strains led to the absence of competition (Fig. 5B). Polar mutations, which can disrupt the expression of downstream genes in an operon, were used alongside non-polar deletions that leave downstream transcription intact, to differentiate the contribution of individual genes versus entire operon disruption. Interestingly, co-culture with the ΔpqsH or ΔpqsL strains resulted in partial reduction of competition similar to the ΔlasR strain (Fig. 5B), indicating that HQNO and PQS may contribute to competition but are not solely responsible for the antagonistic effects on KP. MvfR positively regulates virulence factors such as hydrogen cyanide, rhamnolipids, and phenazines through the activation of PqsE [50–57]. However, co-culturing with ΔpqsE mutant showed similar reductions in viable KP cells as the WT strain (Fig. 5B), suggesting that individual PqsE-regulated molecules are not essential for competition under these conditions.

The relevance of pqsABCDE in competition may stem from the production of antagonistic small molecules or through HHQ/PQS-mediated activation of MvfR, which further induces other antagonistic factors like phenazines [14], siderophores [35], and the type VI secretion system machinery [58]. To better understand the relative contributions of PqsABCDE and MvfR and begin to assess these two possibilities, we complemented the ΔpqsA polar mutant with a plasmid constitutively expressing MvfR (pDN18-mvfR) and ΔmvfR mutant with a plasmid expressing constitutive PqsABCDE (pDN18-pqsABCDE). The results showed that in the absence of MvfR, constitutive expression of pqsABCDE could partially rescue competition with KP (Fig. 5B). By contrast, constitutive expression of mvfR did not rescue the loss of the pqsA (Fig. 5B). These findings imply that MvfR controls competitive interactions with KP by activating the pqs biosynthetic pathway, and while MvfR alone is not sufficient for competition without the pqsABCDE operon, it is crucial for achieving maximal KP killing.

We then tested additional mutants to assess the involvement of other MvfR-regulated antagonistic factors in competition with KP. Neither phenazines nor the type VI secretion systems were solely responsible for inhibiting KP viability in coculture; however, a mutant lacking both pvdA and pchE genes showed significantly reduced KP killing, and a mutant lacking pvdA, pchE, and both phenazine operons fully restored KP viability (Fig. 5B). These experiments demonstrate the multifactorial nature of bacterial competition, with specific genes (e.g., phz1/2) being conditionally relevant depending on the expression level of other genes. Overall, our findings support a model in which PA kills KP through the production of multiple antagonistic factors, primarily involving the Pqs biosynthesis pathway and possibly synergizing with other MvfR-regulated mechanisms.

### 3.6. The PA MvfR-inhibitor D88 attenuates virulence in KP and SA, indicating broad anti-virulence activity

PA, KP, and SA are frequent colonizers of human lungs, causing major health implications. Since an anti-virulence approach does not affect viability, next, we assessed whether D88 treatment attenuates virulence in KP by measuring the biofilm formation and cell adherence of PA and KP in both single- and co-infection settings with or without exposure to D88. First, 50 µM of D88 reduced approximately 40% of biofilm formation in PA following 15 hours of monoculture (Fig 6A). Strikingly, the biofilm formation in KP was also severely impaired by 50 µM of D88, leading to a reduction as high as 73%. Further, biofilm formation was also decreased during the co-culture with PA, revealing the anti-virulent efficacy of D88 against dual-species communities (Fig. 6A). While crystal violet staining quantifies total biomass, it does not distinguish species or separate bacterial load from matrix production. Therefore, we cannot determine whether these reductions stem from one or both species, or from decreased attachment versus matrix disruption. Future experiments will address these distinctions.

**Figure 6.**
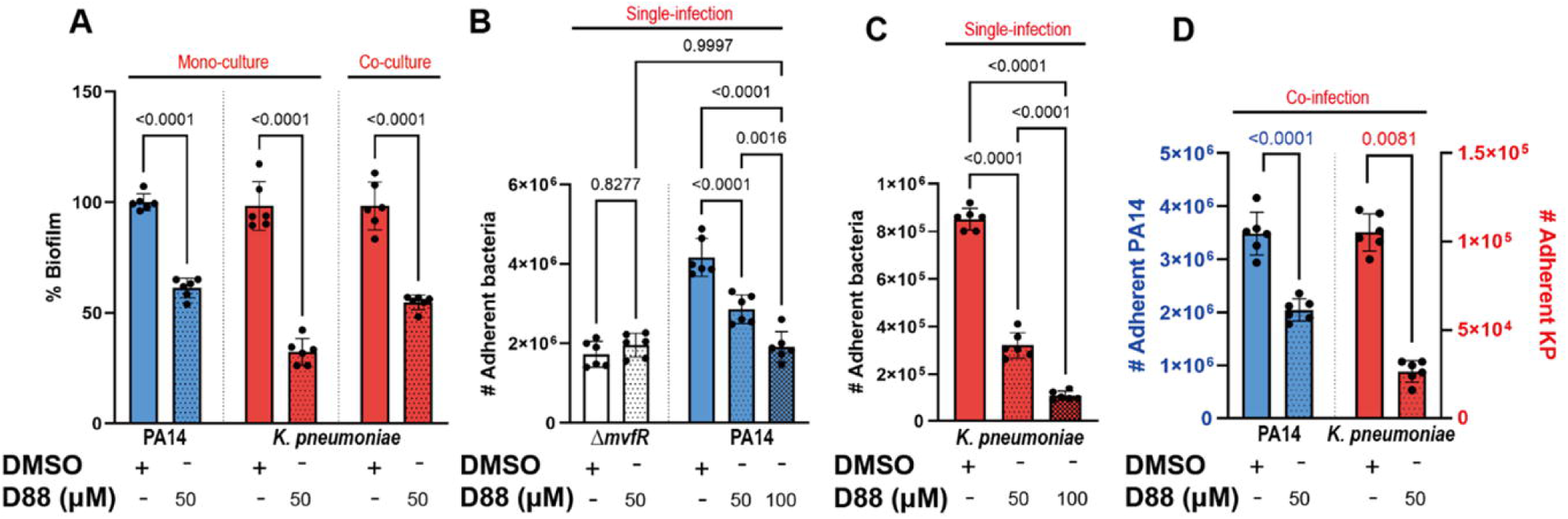
The MvfR-inhibitor (D88) attenuates biofilm formation and adherence to human lung epithelial cells (A549) in PA and KP in single- and co-infection settings. A) Treatment with 50 µM of D88 leads to significant reduction in biofilm formation in PA and KP during mono- and co-culture. Total of 8 × 106 CFU were inoculated for 15 h stationary incubation at 37oC, and the biomass of biofilm was quantified by crystal violet staining. B) Adherence to human lung epithelial cells (A549) and the effect of D88 on adherence was analyzed in P. aeruginosa (PA14), isogenic mutant lacking mvfR, C) KP in a single-infection model. D) Treatment with 50 µM of D88 results in similar decrease in adherence to A549 cells during co-infection with 1:1 ratio of PA and KP. A549 cells at 100% confluency were infected at multiplicity of infection (MOI) of 25:1, and cells were incubated 1.5 h for adherence. The adherent bacteria were differentially enumerated after serial dilution. p-values of one-way ANOVA Šídák’s multiple comparisons test are shown numerically to indicate statistical significance. Complete statistical analysis is presented in Table S5.

To test whether the D88 treatment leads to attenuation in human lung epithelial cell colonization, we adopted the cell adherence assay utilizing the human lung epithelial (A549) cells. Treatment with D88 led to a dose-dependent attenuation in PA adherence to A549 cells, with 100 µM of D88 resulting in a 50% reduction in adherence, which resembled that of a ΔmvfR mutant (Fig. 6B). Notably, D88 also demonstrated a strong attenuating effect in KP adherence to lung epithelial cells, reducing adherence by more than 75% upon exposure to 100 µM (Fig. 6C). During co-infection, the adherence of KP to lung epithelial cells was significantly lower compared to single-infection in the absence of D88 treatment, indicating that KP adherence to A549 cells is profoundly influenced by PA in shared niches (Fig. 6D).

With exposure to 50 µM D88, KP adherence was decreased by approximately 73%. The successful inhibition of MvfR, the lack of selective pressure exerted, and strong attenuation in adherence to both abiotic and biotic surfaces by D88 are strong indicators of a promising candidate for an anti-virulence agent.

Having observed the therapeutic potential of the D88 in a Gram-negative pathogen, we tested whether this inhibitor attenuates the biofilm formation and cell adherence to A549 of a multidrug-resistant Gram-positive that infects many of the same body sites as PA, but unlike KP, was not outcompeted by PA WT in EMEM. Using SA strain USA300, we discovered that 50 µM D88 was sufficient to decrease the SA biofilm formation by approximately 50% over 15 hours of monoculture and by 30% during co-culture with PA14 (Fig. 7A). Adherence to A549 cells was also attenuated in SA by D88 in both single- (Fig. 7B) and co-infection settings (Fig. 7C), as the recovery of cell-bound bacteria after co-infection with PA14 in the presence of 50 µM D88 was significantly lower for both PA14 and SA compared to DMSO treatment. As expected, this adherence assay also revealed a low recovery of ΔmvfR mutant after coinfection with SA in the absence of D88 compared to PA14 (Fig. 7C). Further, the low recovery of cell-bound ΔmvfR after coinfection with SA resembled that of PA14 WT in presence of D88, indicative of the inhibitory action of D88 on MvfR. Similarly, the yield of cell-bound SA after coinfection with ΔmvfR mutant was significantly higher compared to the coinfection with PA14 WT in the absence of D88, underscoring the role of MvfR in competition against the Gram-positive pathogen, SA (Fig. 7). Taken together, these findings suggest that the PA MvfR-inhibitor D88 has broad anti-virulence activity in diverse bacteria, including KP and SA.

**Figure 7.**
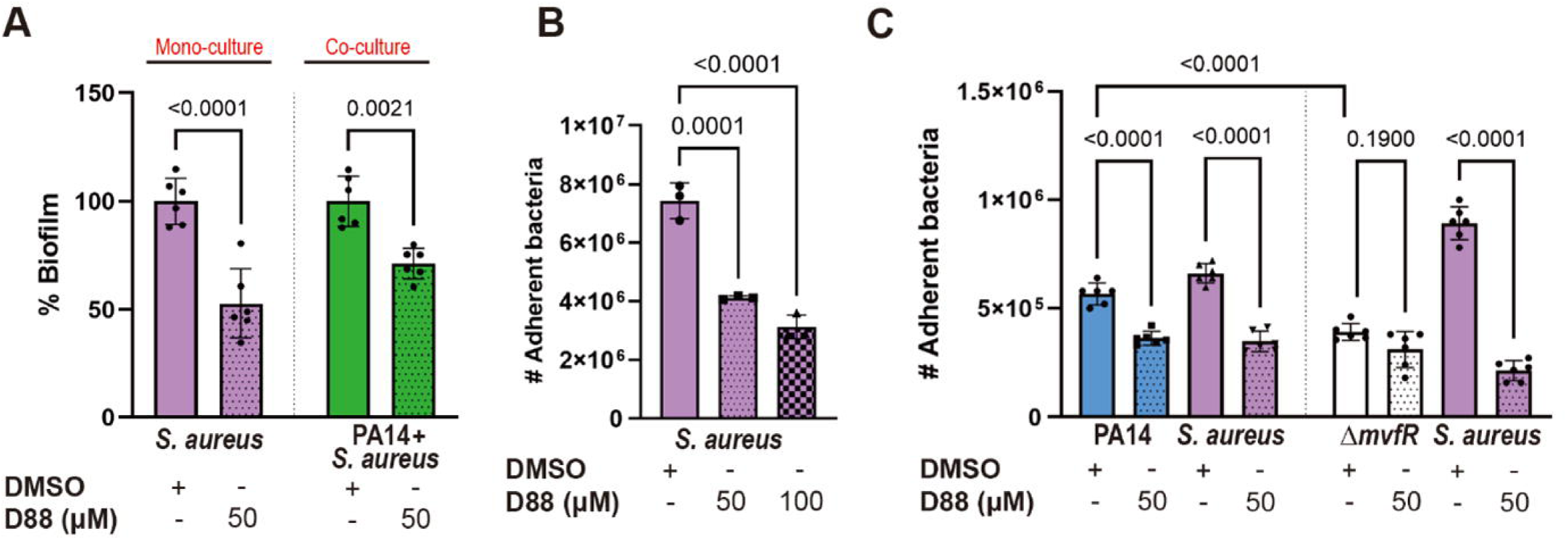
The MvfR-inhibitor D88 significantly diminishes the virulence capacity of the Gram-positive pathogen S. aureus. A) The % biofilm production in SA strain USA300 is significantly impaired upon treatment with 50 µM of D88 during mono- and co-culture. Total of 5 × 106 CFU were inoculated for 15 h stationary incubation at 37oC, and the biomass of biofilm was quantified by crystal violet staining. B) D88 has an inhibitory effect in cell adherence to A549 cells during SA single-infection. C) Treatment with 50 µM of D88 leads to decreased adherence to A549 cells during co- infection with 1:1 ratio of PA14 and SA. A549 cells at 100% confluency were infected at multiplicity of infection (MOI) of 25:1, and cells were incubated 1.5 h for adherence. The adherent bacteria were differentially enumerated after serial dilution. p-values of one-way ANOVA Tukey’s multiple comparisons test are shown numerically to indicate statistical significance. Complete statistical analysis is presented in Table S5.

## 4. Discussion

Polymicrobial infections are exceedingly common in the clinic. This study aimed to identify interactions between some ESKAPEE microbes in polymicrobial communities. In particular, we focused on the interspecies interaction between other important pathogens with PA, which are frequently isolated from polymicrobial infections. By investigating interspecies interactions within defined polymicrobial communities, we can build a framework for understanding which pathogens may suppress the colonization of other microbes during infections and which pathogens may infect the host simultaneously.

Using a dual-species in vitro culturing system and a library of PA QS mutants, we showed that two-member microbial communities engage in complex interactions, including interference competition with other ESKAPE Gram-negative bacteria, but not with the Gram-positive pathogens. Specifically, we found that PA and SA coexist in static EMEM culture for at least 24 h. From a technical standpoint, this is an interesting finding because SA and PA often coexist in infections, but PA quickly outcompetes SA in most in vitro settings [59, 60].

Importantly, we discovered that among the three major quorum-sensing regulators, MvfR, encoded by the multiple virulence factor regulator gene (mvfR), is necessary for competition with each of the Gram-negative competitors, including KP, AB, and EC. This finding suggests that MvfR-dependent competition may be a general strategy evolved to compete with pathogenic and commensal microbes in diverse environments and gain a competitive foothold in these niches, potentially resulting in dysbiosis. MvfR controls the production of numerous antagonistic factors [17, 44], including siderophores [35], HQNO [24, 49], PQS [16], phenazines [14], and the type VI secretion system machinery [58], which likely aid PA in competing with diverse, especially Gram-negative microbes. The role of the MvfR/PqsABCDE system in interspecies competition has been well established in previous studies [61–63], which demonstrated its involvement in toxin regulation and virulence. Our findings build upon this foundation by further elucidating MvfR’s specific contribution to competition with ESKAPEE pathogens.

Compared to monomicrobial infections, polymicrobial infections are challenging to treat for several reasons. One potential consequence of a polymicrobial infection is that interactions between competing microbes commonly impact antibiotic susceptibility relative to a monomicrobial infection. Depending on which microbes are present, broad-spectrum or combinatorial antibiotics may be required, but this approach presents a few dilemmas. First, broad-spectrum antibiotics cannot differentiate between pathogens and commensals, disrupting the microbial flora and increasing the risk of secondary infection [64]. Second, antibiotics create a selective pressure that drives the evolution of resistance, making infections recalcitrant to conventional therapies and leading to high mortality. Manipulating bacterial virulence to aid in the treatment of infection rather than killing pathogens is an appealing approach that has been successfully applied to burn and lung infections in mice [30, 34, 65].

Here, we examine how the potent anti-virulence agent, D88, an N-Aryl Malonamide that inhibits MvfR, alters the composition of polymicrobial communities. Although D88 reduces competition between PA and other ESKAPEE pathogens, permitting their viability, it disarms them, decreasing their virulence. Thus, the possibility of promoting viability in a polymicrobial infection doesn’t exclude the possibility of reduced infectivity when applying an anti-virulence approach to treat some infections. This effect would be highly desirable for reducing competition in the context of dysbiosis, as it would allow for commensal communities to remain intact during treatment. Studying how anti-virulence therapies influence these more complex microbiota that include beneficial microbial species will be an important area of future research.

Given the ability of D88 to reduce biofilm formation and host cell attachment in both KP and SA, two phylogenetically distant bacteria, it will be exciting to investigate what this inhibitor targets. MvfR is a LysR-type transcriptional regulator (LTTR), the most common type of prokaryotic DNA-binding protein. It is possible that while D88 targets MvfR in PA, it may also target another LTTR that controls functions related to microbial competition, host cell and abiotic surface attachment, or virulence in other bacteria. For instance, in KP the LysR-type regulator OxyR is involved in mucosal and abiotic colonization [66] and therefore may be a putative target of D88 and responsible for the observed reduction in surface attachment. Likewise, SA USA300 encodes at least seven LTTRs [67] that could serve as potential targets for D88. Of particular interest is the regulator LTTR852, which is crucial for secondary tissue colonization [68]. However, it is also possible that D88 targets a non-LTTR protein involved in bacterial competition. Whether D88 effects on co-cultures stem solely from reduced virulence or also involve broader metabolic changes in either PA or other microbes is of particular interest and warrants further investigation.

While D88 increases KP survival in planktonic co-culture by inhibiting PA QS-dependent antagonism, it reduces KP adherence in the biofilm context. This suggests that D88 may exert context-dependent effects—reducing *P. aeruginosa*- mediated killing while also modulating KP surface colonization. By attenuating virulence traits like adherence without impacting viability, D88 may influence interspecies dynamics and host interactions in a way that minimizes selective pressure and may not carry the same risk of resistance development as traditional antibiotics.

Overall, this study provides a framework for studying the impact of QS and anti-virulence therapy on interactions within polymicrobial communities during infection and how community structure can be manipulated with anti-virulence therapeutics. The environmental context is likely an important determinant of the outcome of bacterial competition since metabolism, growth phase, and bacterial density can each influence the transcriptional profiles and competitiveness of PA and its competitors. The knowledge obtained from this study can provide fundamental insights into microbial interactions, which have broad implications not only for treating acute and chronic infections, including in the lungs of people with cystic fibrosis, and in burn and diabetic wounds. Our findings highlight the pivotal role of MvfR in controlling competition and virulence, and we anticipate that anti-virulence drugs, like the MvfR inhibitor D88, may hold promise for restoring microbial homeostasis in settings where PA antagonizes commensal microbes. However, successful implementation requires careful consideration of the infection context, especially in infection environments like the lung or an injury site where low oxygen or static conditions may prevent a quorum from being reached, or where PA might suppress other opportunistic microbes. In this scenario, the application of an anti-virulence compound might be beneficial or deleterious depending on the anti-virulence compound used and its impact on a polymicrobial infection. Additionally, comparative studies directly benchmarking the efficacy, safety, and resistance development of anti-virulence therapies to conventional antibiotics are needed to fully assess their therapeutic potential.

The results of this study may also provide insights into interactions occurring outside the infection setting where these bacteria share other ecological niches. For example, PA, KP, and SA are ubiquitous microbes naturally occurring in the environment, including in soil, plant-associated niches, and water. Understanding how QS systems impact microbial interactions in these contexts may set the stage for developing anti-virulence strategies with direct non-human applications, such as veterinary, agricultural, and industrial uses. Future studies should assess bacterial competitive and cooperative interactions in the presence and absence of therapeutics in physiologically relevant environments that closely mimic human infections. While our in vitro dual-species co-culture system provides valuable mechanistic insights, incorporating animal models or clinical samples will be crucial for validating these findings and further understanding the role of QS in microbial competition within host environments.

## 5. Conclusions

Our findings identify the quorum-sensing regulator MvfR as a key mediator of interspecies competition by PA against other Gram-negative ESKAPEE pathogens, including KP, AB, and EC. Using a defined dual-species co-culture system, we show that MvfR-dependent interference competition contributes to community composition and PA dominance. Importantly, inhibition of MvfR with the anti-virulence compound D88 attenuates virulence and reshapes microbial interactions without affecting bacterial viability. These results provide mechanistic insights into QS-driven interbacterial dynamics and support further investigation into anti-virulence therapies as microbiota-sparing alternatives to conventional antibiotics. Future work should evaluate these effects in more physiologically relevant models to assess translational potential.

## Contributions

L.G.R. and K.M.W. contributed to the conception of the study. K.M.W., M.W.O. and L.G.R. contributed to the design of the study and wrote the paper. K.M.W., M.W.O., J.F., E.K., and A.D. completed competition experiments. M.W.O. and A.M. completed biofilm assays. M.W.O completed A459 adherence experiments. All authors contributed to data analysis and interpretation. All authors have approved the submitted version of the study and their contributions.

## Acknowledgments

This study was supported by the Massachusetts General Hospital Research Scholar Award and grant R01AI177555 to LGR. We thank Enamine Ltd., Kyiv, Ukraine for D88 synthesis. The funders had no role in study design, data collection, interpretation, or the decision to submit the work for publication. We acknowledge the Glue Grant (“Inflammation and Host Response to Injury”) Consortium for the dataset we analyzed.

## Data Availability Statement

All data generated during the current study are available from the corresponding author upon reasonable request. Availability to the Glue grant patient data will require an IRB.

