## supplemental data for "MvfR shapes *Pseudomonas aeruginosa* Interactions in Polymicrobial Contexts: Implications for Targeted Quorum Sensing Inhibition"

**Content:**

Figure S1-S6

List of Supplementary Tables

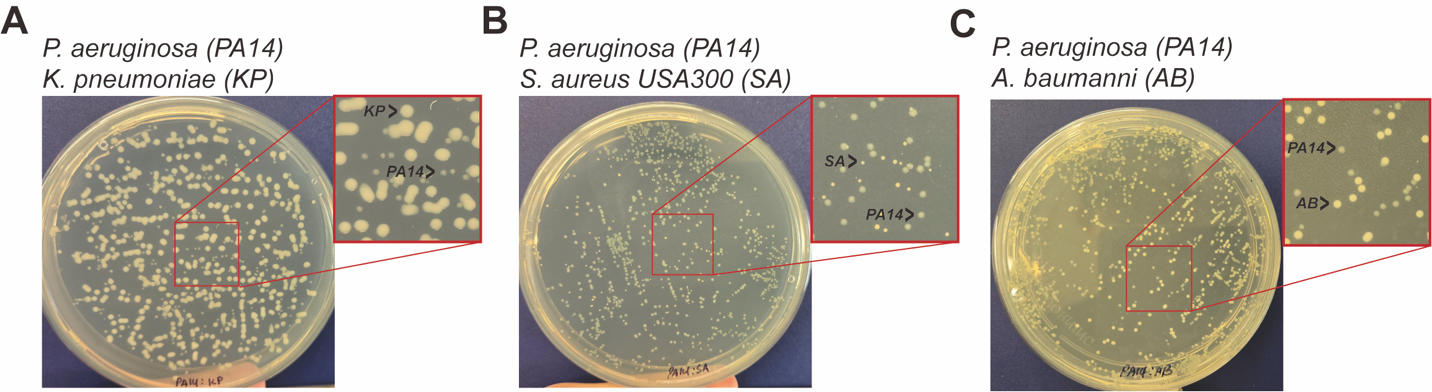

**Figure S1. *P. aeruginosa* (*PA14*) can be differentially enumerated on LB agar plate based on colony morphology after coculture with either *K. pneumoniae* (*KP*), *S. aureus* USA300 (*SA*), or *A. baumanni* (*AB*)***. PA14* was cocultured with either **A)** *KP,* **B)** *SA,* or **C)** *AB* overnight at 37 °C on an LB agar plate. The unique colony morphology of each species allows for differential enumerations on non-selective LB agar plates.

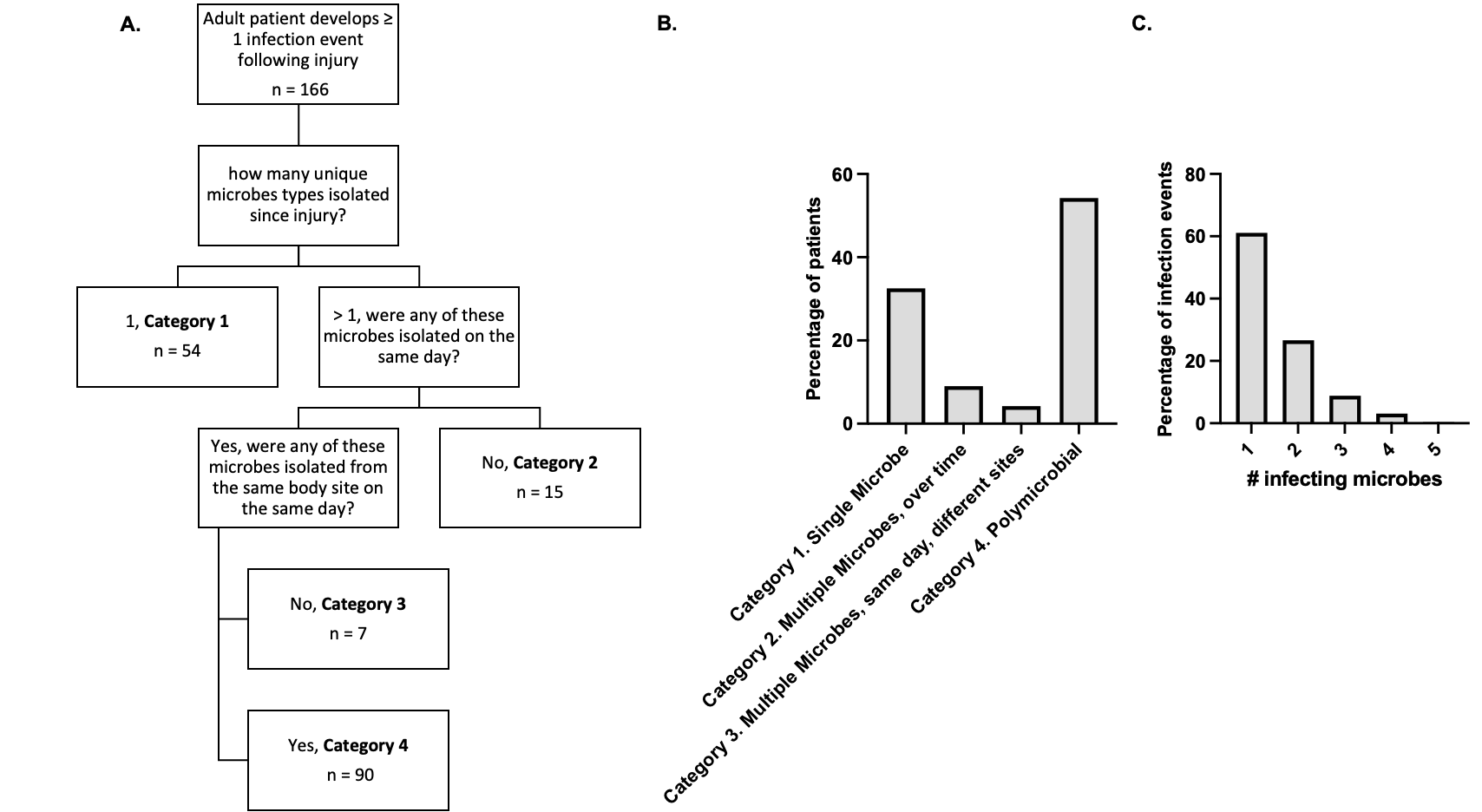

**Figure S2. Analysis of the Glue Grant.** (A) Decision tree for classification of patients as: Category 1. infected by a single microbe type across time (Single Microbe), Category 2. multiple infections over time involving different types of microbes (Multiple Microbes, over time), Category 3. Multiple types of monomicrobial infection across body site (Multiple Microbes, same day, different sites), and Category 4. at least 1 infection with multiple infecting microbes at a single body site on a single day (Polymicrobial). (B) The proportion of patients in the Glue Grant study falling into each category. (C) The proportion of infection events (i.e., isolation of ≥ 1 microbe isolated from a particular body site per day) involving 1 – 5 microbe types. Microbial counts are presented in Table S1.

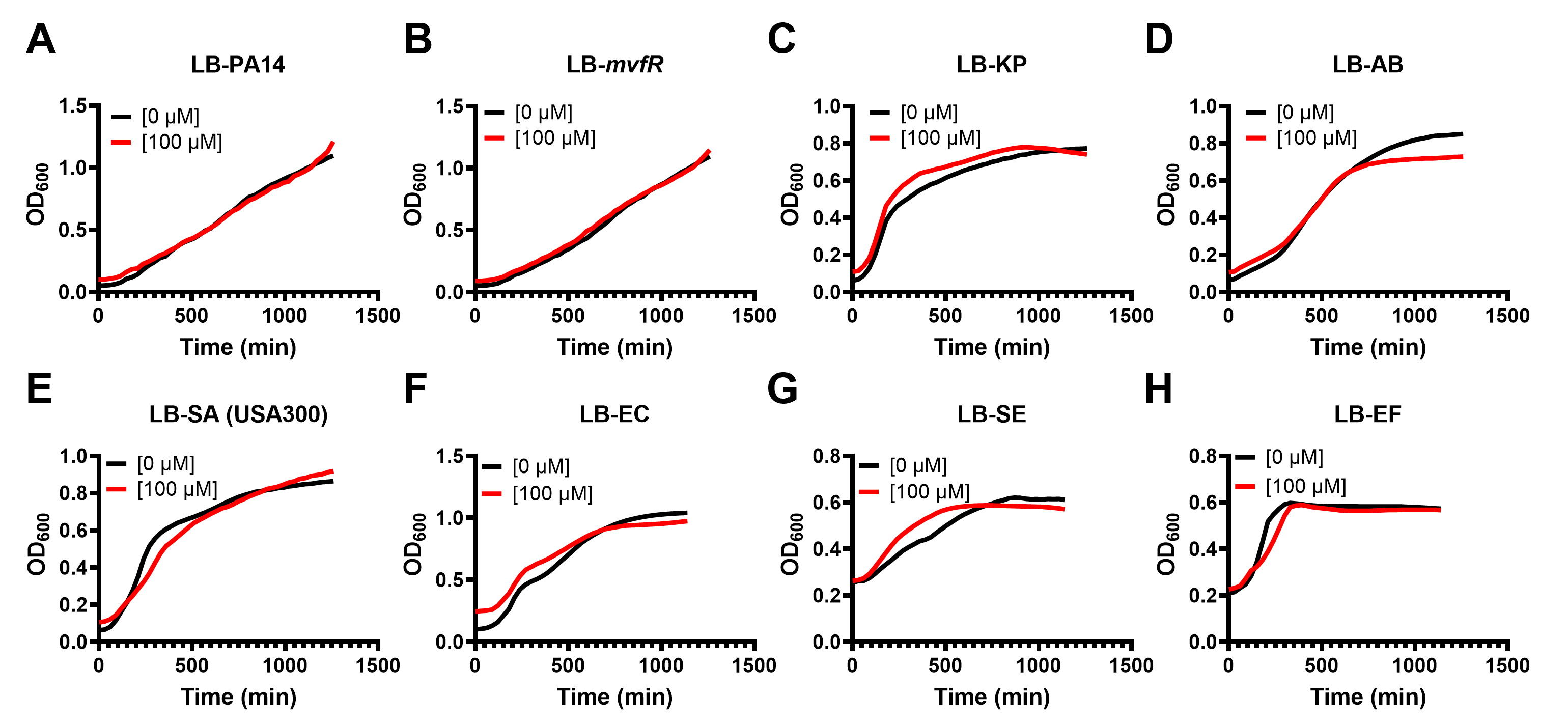

**Figure S3. Growth kinetics of bacterial species in the presence of D88 in Luria Broth (LB).** 100 µM of D88 has negligible impact on the growth of A) PA14, B) PA14 ∆*mvfR*, C) *K. pneumoniae,* D) A*. baumannii,* E) *S. aureus* USA300, F) *E. coli,* G) *S. epidermis,* or H) *E.* *faecalis* in LB.

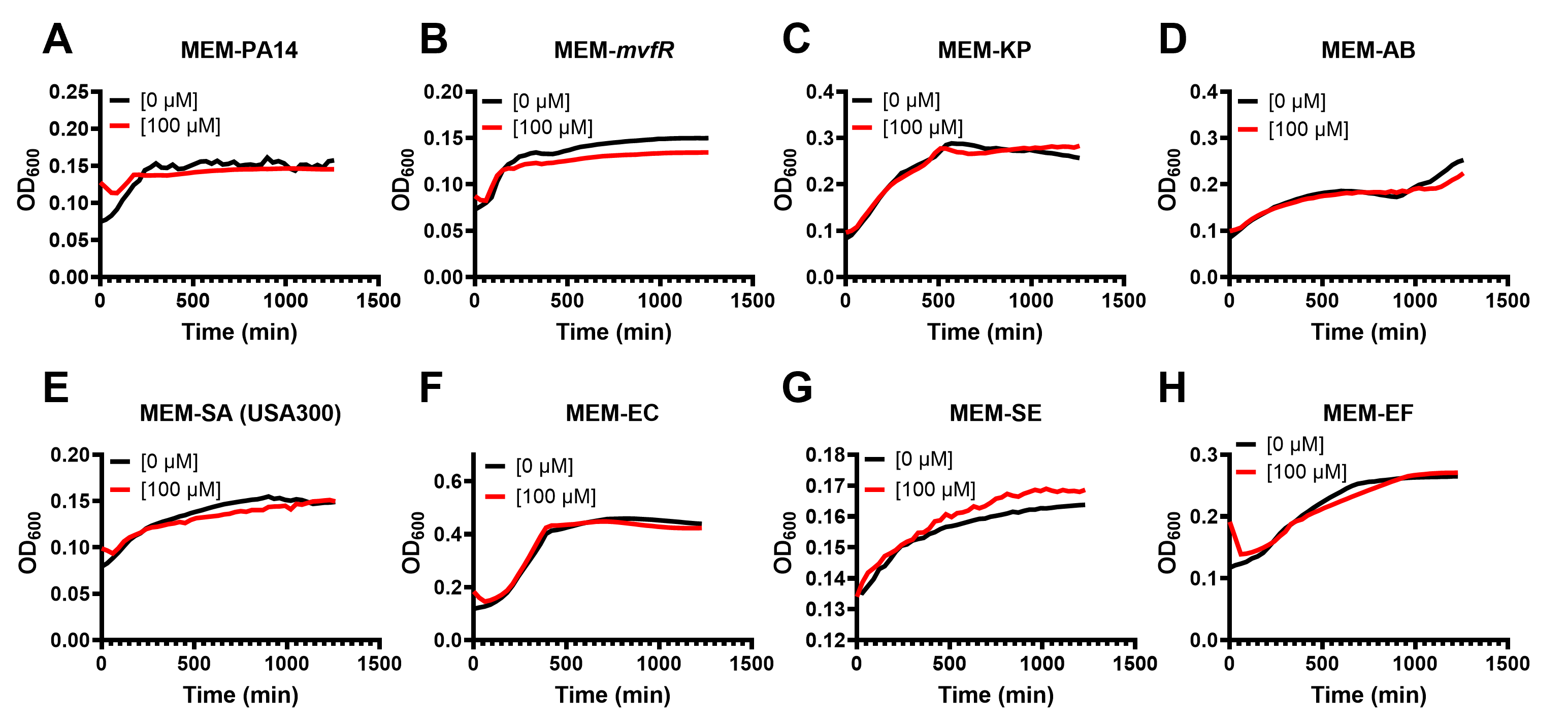

**Figure S4. Growth kinetics of bacterial species in presence of D88 in Minimum Essential Medium (MEM).** 100 µM of D88 has negligible impact on the growth of A) PA14, B) PA14 *∆mvfR,* C) *K. pneumoniae,* D) *A. baumannii,* E) *S. aureus* USA300, F) *E. coli,* G) *S. epidermis,* or H) *E. faecalis* in MEM*.*

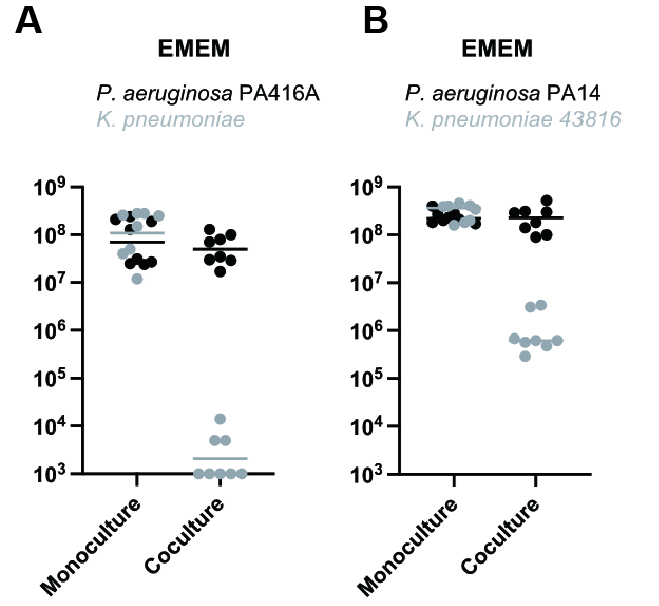

**Figure S5. Competition between different isolates of *P. aeruginosa* and *K. pneumoniae.* (A)** Enumeration of *P. aeruginosa* PA416A and *K. pneumoniae* CFU following 24 h growth in monoculture or co-culture in EMEM. *P. aeruginosa* CFU for each replicate is indicated with black circles, and *K. pneumoniae* CFU for each replicate is indicated with grey circles, *n = 8* biological replicates. The middle line indicates the median log-transformed CFU/mL per bacterium in monoculture or co-culture**. (B)** Enumeration of *P. aeruginosa* PA14 and *K. pneumoniae* 43816 CFU following 24 h growth in monoculture or co-culture in EMEM. *P. aeruginosa* CFU for each replicate is indicated with black circles, and *K. pneumoniae* CFU for each replicate is indicated with grey circles, *n = 8* biological replicates. The middle line indicates the median log-transformed CFU/mL per bacterium in monoculture or co-culture.

For all panels, inoculating CFU at 0h was 0.5-1 x 10^7^ CFU/mL. Significance was assessed by two-way ANOVA on log_10_-transformed data with Šídák's multiple comparisons test. Statistical analysis is presented in Table S7.

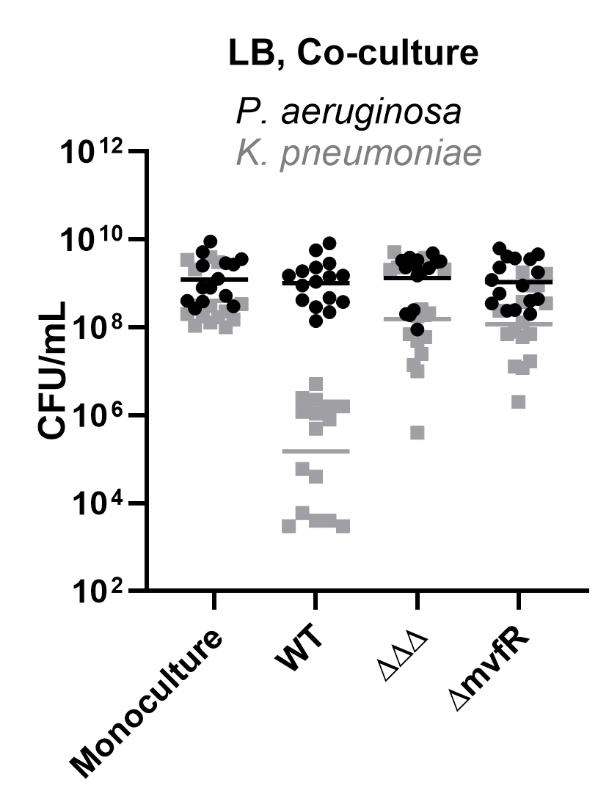

**Figure S6. Enumeration of *P. aeruginosa* WT, ∆*mvfR*, or the LasR, RhlR, MvfR triple mutant (∆∆∆), and *K. pneumoniae* CFU following 24 h growth in monoculture or co-culture in LB.** *P. aeruginosa* CFU for each replicate is indicated with black circles, and *K. pneumoniae* CFU for each replicate is indicated with grey squares, *n = 16* biological replicates. The middle line indicates the median log-transformed CFU/mL per bacterium in monoculture or co-culture. Inoculating CFU at 0h was 0.5-1 x 10^7^ CFU/mL. Significance was assessed by two-way ANOVA on log_10_-transformed data with Dunnett’s multiple comparisons test. Statistical analysis is presented in Table S8.

**List of Supplementary Tables**

**Table S1.** Counts, percentages, and ranks of infecting organisms in the Glue Grant dataset.

**Table S2**. Statistical analysis of data in Figure 2: Two-way ANOVA on log_10_-transformed data, Šídák's multiple comparisons test. Significant differences with a |(mean log10-transformed change in abundance)| > 0.5 are bolded.

**Table S3**. Statistical analysis of data in Figure 3: Two-way ANOVA on log_10_-transformed data, Dunnett’s multiple comparisons test. Significant differences with a |(mean log10-transformed change in abundance)| > 0.5 are bolded.

**Table S4.** Statistical analysis of data in Figure 5: Two-way ANOVA on log_10_-transformed data, Dunnett’s multiple comparisons test. Significant differences with a |(mean log10-transformed change in abundance)| > 0.5 are bolded.

**Table S5.** Statistical analysis of data in Figure 6. Statistical significance p < 0.05 are bolded.

**Table S6**. Statistical analysis of data in Figure 7. Statistical significant p<0.05 are bolded.

**Table S7**. Statistical analysis of data in Figure S5: Two-way ANOVA on log_10_-transformed data, Šídák's multiple comparisons test. Significant differences with a |(mean log10-transformed change in abundance)| > 0.5 are bolded.

**Table S8**. Statistical analysis of data in Figure S6: Two-way ANOVA on log_10_-transformed data, Dunnett’s multiple comparisons test. Significant differences with a |(mean log10-transformed change in abundance)| > 0.5 are bolded.
